# Interactions across life stages and thermal tolerance plasticity in *Tigriopus californicus*

**DOI:** 10.1101/751040

**Authors:** Timothy M. Healy, Antonia K. Bock, Ronald S. Burton

## Abstract

In response to rapid environmental change, organisms rely on both genetic adaptation and phenotypic plasticity to adjust key traits that are necessary for survival and reproduction. Given the accelerating rate of climate change, plasticity may be particularly important. For organisms in warming aquatic habitats, upper thermal tolerance is likely to be a key trait, and many organisms express plasticity in this trait in response to developmental or adulthood temperatures. Although plasticity at one life stage may influence plasticity at another life stage, relatively little is known about these interactive effects for thermal tolerance. Here we used locally adapted populations of the intertidal copepod *Tigriopus californicus* to investigate these interactions in a marine ectotherm. We found that low latitude populations had greater critical thermal maxima (CT_max_) than high latitude populations, and variation in developmental temperature altered CT_max_ plasticity in adulthood. After development at 25°C, CT_max_ was plastic in adults, whereas no adult plasticity in this trait was observed after 20°C development. This pattern was identical across four populations, suggesting that local thermal adaptation has not shaped this interactive effect. However, differences in the capacities to maintain ATP synthesis rates and to induce heat shock proteins at high temperatures, two likely mechanisms of local adaptation in this species, were consistent with changes in CT_max_ due to developmental temperatures, suggesting there is mechanistic overlap between plastic interactions and adaptation in general. These results indicate that interactive effects of plasticity across life stages may have substantial impacts on upper thermal tolerance in ectothermic organisms.

**Summary statement:** Developmental temperatures alter the plasticity of thermal limits in adults of a marine ectotherm, and differences in ATP synthesis rate and heat shock protein expression parallel the changes in tolerance.

## Introduction

As the earth warms, organisms are increasingly impacted by the effects of high environmental temperatures (e.g., Wiens, 2016; Cohen et al., 2018; Pinsky et al., 2019). Indeed, the geographic range limits of many species have already shifted as a result of anthropogenic climate change, and in general these shifts have been towards the poles (i.e., to cooler temperatures; e.g., Parmesan and Yohe, 2003; Perry et al., 2005; Chen et al., 2011). The extents to which these effects have occurred, and will continue to occur, depend largely on the adaptive and plastic capacities of organisms to adjust key physiological traits, such as thermal tolerance limits (especially in aquatic ectotherms; Sunday et al., 2012; Pinsky et al., 2019), in response to increased temperatures (e.g., Crain et al., 2008; Somero, 2010; Bay et al., 2017; Kellermann and van Heerwaarden, 2019). In particular, given that rapid phenotypic changes are necessary due to high rates of climate change (e.g., Barrett and Hendry, 2012; Fox et al., 2019), phenotypic plasticity may play a critical role in the resilience of populations and species to the effects of climate change (Merilä and Hendry, 2014; Seebacher et al., 2015; Donelson et al., 2019; Morley et al., 2019).

Phenotypic plasticity occurs across life stages and generations (e.g., Kelly et al., 2011; Schulte et al., 2011; Beaman et al., 2016; Burggren, 2015). For example, temperatures experienced during development or adulthood often have irreversible or reversible effects on physiological traits (e.g., Schulte et al., 2011; Beaman et al., 2016), and multi- or trans-generational effects of thermal variation are commonly observed (e.g., Crill et al., 1996; Massamba-N’Siala et al., 2014; Zizzari and Ellers, 2014; Donelson et al., 2018). Thus, physiological phenotypes have the potential to be shaped by interactive effects of plasticity across different life stages. However, compared to interactions between adaptive processes and phenotypic plasticity (e.g., Crispo, 2007; Hendry, 2016; Donelson et al., 2019; Kelly, 2019), effects of plasticity at one life stage on the expression of plasticity at another life stage have received relatively little attention (Beaman et al., 2016). That said, interactions across life stages have been observed for several traits (reviewed in Beaman et al., 2016). For instance, differences in developmental conditions alter the phenotypic plasticity of swimming performance and metabolic rate in adult mosquitofish (*Gambusia holbrooki*; Seebacher et al., 2014; Seebacher and Grigaltchik, 2015). Yet, despite the likely biogeographic importance of thermal tolerance limits (Sunday et al., 2012), and many published examples of thermal tolerance limit plasticity in ectothermic organisms as a result of developmental or adulthood temperatures (e.g., Stillman and Somero, 2000; Ford and Beitinger, 2005; Fangue et al., 2006; Angiletta, 2009; Overgaard et al., 2011; Cooper et al., 2012; Tepolt and Somero, 2014; Jakobs et al., 2015; Troia et al., 2015; Kingsolver et al., 2016; Pereira et al., 2017; Diamond et al., 2018; Mueller et al., 2019; Yanar et al., 2019), relatively few studies have assessed the potential for interactions between these life stages to shape the phenotypic plasticity of upper thermal tolerance (although see Schaefer and Ryan, 2006; Kellermann et al., 2017; Kellermann and Sgrò, 2018). Here we examine the effects of developmental temperature on the expression of upper thermal limit plasticity, and the potential mechanistic basis for these effects in populations of the intertidal copepod *Tigriopus californicus*.

*T. californicus* are small (∼1.2 mm) harpacticoid copepods with short generation times (3-4 weeks) that inhabit supralittoral tidepools along the west coast of North America from Baja California, Mexico to southern Alaska, USA. Populations of this species occur on rocky outcrops isolated by sandy beaches with very low gene flow and high levels of genetic divergence among populations (Burton and Lee, 1994; Burton, 1997, 1998; Edmands, 2001; Peterson et al., 2013; Pereira et al., 2016; Barreto et al., 2018). Although much of this genetic divergence is likely a result of small effective population sizes and genetic drift acting on selectively neutral variation, signatures of directional selection have been detected across the transcriptome (Pereira et al., 2016) indicating that at least a portion of the genetic differentiation among populations is likely adaptive. Moreover, several common-garden studies in laboratory-raised individuals have demonstrated differences in upper and lower thermal tolerance limits that are consistent with local thermal adaptation in response to the latitudinal temperature gradient across the species range (Willett, 2010; Kelly et al., 2012; Wallace et al., 2014; Pereira et al., 2014, 2017; Leong et al., 2018; Willett and Son, 2018; Foley et al., 2019). This variation among populations has also been associated with genetically based variation in the function and regulation of heat shock protein genes (Schoville et al., 2012; Barreto et al., 2015; Tangwancharoen et al., 2018) and in the maintenance of mitochondrial ATP synthesis rates at high temperatures (Harada et al., 2019). Few studies have examined temperature-mediated phenotypic plasticity in these traits in *T. californicus*, but elevated developmental temperature is known to increase upper thermal tolerance regardless of population (Kelly et al., 2012, 2017; Pereira et al., 2017), whereas adult plasticity in this trait is thought to be limited (although only relatively short acclimation periods have been examined [e.g., 1 d]; Pereira et al., 2017). Taken together with short generations and ease of laboratory culture, these observations make *T. calfornicus* an ideal study system in which to investigate interactive effects of plasticity at different life stages in an aquatic ectotherm.

In the current study, we use laboratory-raised *T. californicus* to test two hypotheses: (1) variation in developmental temperatures changes the expression of phenotypic plasticity of upper thermal tolerance in adults, and (2) the physiological mechanisms involved in local thermal adaptation among populations are also involved in thermal limit plasticity associated with temperatures experienced during development. First, we expand and validate our previous study (Harada et al., 2019) establishing critical thermal maximum (CT_max_) methods to determine upper thermal tolerance in this species. We then use this method (which reduces the sample sizes required to estimate upper thermal tolerance) to facilitate split-clutch experiments examining interactive effects of developmental and adulthood temperatures on upper thermal tolerance in four Californian populations of *T. californicus*. Finally, we assess the effects of developmental temperature on traits involved in local thermal adaptation: the thermal performance curve of ATP synthesis rate, and the mRNA expression levels of heat shock protein genes and mitochondrial-encoded genes following acute heat stress.

## Materials and methods

### Collection and culturing of copepods

Adult copepods were collected from supralittoral tidepools across ten locations along the west coast of North America, which spanned ∼21.5° of latitude (San Roque, Mexico – SR, La Bufadora, Mexico – BF, San Diego, California – SD, Bird Rock, California – BR, Abalone Cove, California – AB, Estero Bay, California – EB, San Simeon, California – SS, Santa Cruz, California – SC, Pescadero, California – PE, and Pacific Crest, Canada – PC; Table S1; Fig. S1A,B). Collected animals were transported to Scripps Institution of Oceanography (San Diego, CA) in 1 L plastic bottles containing seawater obtained from the same tidepools. The collection for each location was divided across several laboratory cultures, which were maintained at 20°C, 36 ppt and 12:12 h photoperiod (light:dark) using filtered seawater and deionized water to adjust salinity as necessary. Laboratory cultures were maintained for at least two generations (∼2 months) prior to experiments. During laboratory acclimations and experimental treatments, copepods consumed natural algal growth within the cultures, as well as a mixture of ground Spirulina (Salt Creek, Inc., South Salt Lake City, UT) and TetraMin Tropical Flakes (Spectrum Brands Pet LLC, Blacksburg, VA) that was added approximately once per week.

### Critical thermal maximum variation among populations

Upper thermal tolerance was measured by critical thermal maximum (CT_max_) trials using loss of locomotor performance as the assay endpoint (as in Harada et al., 2019). In brief, sixteen adult copepods of each population (8 females and 8 males for all populations; divided across five trials) were transferred to 10-cm petri dishes containing filtered seawater (20°C and 36 ppt) with no food overnight. In the morning, copepods were individually transferred into 0.2-mL strip tubes with 100 µL of water from the petri dish. Tubes were left uncapped, and were placed in an Applied Biosystems SimpliAmp™ Thermal Cycler (Thermo Fisher Scientific, Waltham, MA). After 5 min at 20°C, the temperature was increased using the AutoDelta function at rates of 0.1°C per 20 s from 20 to 32°C, and 0.1°C per min from 32°C to the temperature at which the last individual in the trial lost locomotor performance. Loss of performance (i.e., knockdown) was monitored by cycling 40 µL of water in each tube with a pipettor. Typically this procedure results in erratic swimming behaviour in *T. californicus*; however, at extremely high temperatures, this swimming response ceases, and copepods passively sink to the bottom of their tube. Endpoints were determined when an individual did not respond to three sequential tests with the pipettor, and CT_max_ was recorded as the temperature at which the endpoint was observed. After CT_max_ was determined, copepods were returned to 10-cm petri dishes with 20°C filtered seawater (36 ppt) for recovery, and survivorship was >90%.

### Developmental and adult plasticity in critical thermal maximum

To assess variation in the phenotypic plasticity of upper thermal tolerance in adult copepods as a result of differences in developmental temperature, gravid SD and BR females with mature (i.e., red) egg sacs were removed from laboratory cultures (24 females from 4 cultures per population). Egg sacs were dissected from the females (which synchronizes hatching), placed individually in wells of 6-well plates containing filtered seawater (20°C and 36 ppt), and allowed to hatch overnight. In the morning, nauplii (i.e., larvae) from each egg sac were counted, and split evenly across six treatments in 10-cm petri dishes. Three treatments were developed at 20°C for 14 d, and three treatments were developed at 25°C for 10 d. Preliminary trials with the SD population determined that these developmental times were those required for the majority of individuals to reach adulthood (and to observe the first gravid female) at each temperature, suggesting that the temperature coefficient (Q_10_) of developmental rate equals ∼2 in this species. At the end of the developmental periods, the developmental treatments at each temperature were transferred to one of three adult acclimations: 20°C for 14 d, 25°C for 10 d, or 25°C for 14 d. These lengths of acclimations were chosen to allow two weeks acclimation at 20°C, and comparisons between equivalent acclimations using either absolute time or physiologically adjusted time at 25°C (assuming a continued Q_10_ of ∼2 for life history traits). On the days that the adult acclimations were completed, critical thermal maxima were determined as described above for 16 copepods from each treatment and population. Note that individuals used in the tolerance trials were transferred to fresh filtered seawater without food at their acclimation conditions on the evening before the end of the acclimation treatments (i.e., the day before trials), and CT_max_ trials started from the acclimation temperatures in all cases.

To examine the potential for local thermal adaptation of the effects of developmental temperature on adult thermal tolerance plasticity, we performed a second experiment beginning with gravid females from the SC and PE populations. This experiment was conducted as described above for the SD and BR populations; however, the 25°C 10 d adult acclimation treatments were excluded, meaning egg sacs for each population were split across four treatments in total (i.e., 20°C development and adulthood; 20°C development and 25°C adulthood; 25°C development and 20°C adulthood; 25°C development and adulthood). Again, CT_max_ was determined for 16 copepods for all treatments except the PE 25°C development and 25°C adulthood treatment for which n = 15.

### Plasticity of ATP synthesis rate thermal sensitivity

To determine phenotypic plasticity of the thermal performance curve for ATP synthesis rate associated with differences in developmental temperature, gravid SD and BR females carrying mature egg sacs (60 per population) were transferred from laboratory cultures to 10-cm petri dishes (6 per population) containing ∼60 mL of filtered seawater (20°C and 36 ppt) with food. The egg sacs from the majority of these females (6-10 per plate) hatched overnight. All females were removed in the morning; egg sacs that were still carried by females were dissected free and returned to their respective dishes. Dissected egg sacs hatched within 3 h, and once all egg sacs had hatched, three petri dishes for each population were transferred to 25°C. As described above, juveniles in the 20 or 25°C dishes developed for 14 or 10 d, respectively. All dishes were kept at their developmental temperatures for adult acclimations, which were also 14 or 10 d at 20 or 25°C, respectively.

ATP synthesis rates were measured at 20, 25, 30, 33, 35 and 37°C using procedures similar to those of Harada et al. (2019). On the day before the end of the adult acclimation treatments, groups of 32 copepods (6 groups per population x treatment) were held at their acclimation temperatures in 10-cm petri dishes with filtered seawater (36 ppt) and no food overnight. In the morning, the groups of copepods were rinsed with 200 µL homogenization buffer (400 mM sucrose, 100 mM KCl, 70 mM HEPES, 6mM EGTA, 3 mM EDTA, 1% w/v BSA, pH 7.6), which had been chilled on ice. Each group was transferred to a 2-mL glass teflon homogenizer, and homogenized in 800 µL of fresh buffer. Following homogenization, mitochondria were isolated by differential centrifugation in 1.5 mL microcentrifuge tubes (Eppendorf, Hamburg, Germany). First, the tubes were centrifuged at 4°C and 1,000 g for 5 min, and supernatants were transferred to fresh tubes. Second, the new tubes were centrifuged at 4°C and 11,000 g for 10 min. The resulting supernatants were discarded, and mitochondrial pellets were resuspended in 205 µL assay buffer (560 mM sucrose, 100 mM KCl, 70 mM HEPES, 10 mM KH_2_PO_4_, pH 7.6). Isolated mitochondria were divided into eight 25-µL aliquots: 6 for synthesis reactions (1 per temperature), 1 for initial ATP concentration determination, and 1 for measuring DNA content which was used to normalize ATP synthesis rate. DNA content was assayed with Invitrogen™ Quant-iT™ PicoGreen™ dsDNA reagent following the manufacturer’s protocols (Thermo Fisher Scientific, Waltham, MA).

ATP synthesis reactions were conducted in 0.2-µL strip tubes, and were initiated by adding 5 µL of a saturating substrate cocktail (final assay substrate concentrations 5 mM pyruvate, 2 mM malate, 10 mM succinate and 1 mM ADP in assay buffer), resulting in election donation to both complex I and complex II (CI+II) of the electron transport system. Following substrate addition, tubes were immediately transferred to an Applied Biosystems SimpliAmp™ Thermal Cycler (Thermo Fisher Scientific, Waltham, MA), and incubated at the desired assays temperatures for 10 min. At the end of the reactions, 25 µL of each assay was added to 25 µL of CellTiter-Glo (Promega, Madison, WI), which stops ATP synthesis, and is used for ATP quantification. To determine initial ATP concentrations in the assays, one aliquot of each mitochondrial isolation was added to CellTiter-Glo immediately following substrate addition. All assays were held in the dark at room temperature for 10 min after addition of CellTiter-Glo, and then luminescence was determined with a Fluoroskan Ascent® FL (Thermo Fisher Scientific, Waltham, MA). ATP concentrations were calculated by comparison to a prepared standard curve (5 nM to 10 µM in assay buffer), and synthesis rates at each temperature were determined by subtracting the initial ATP concentration from the final ATP concentrations at each temperature for each mitochondrial isolation.

### Plasticity of gene expression following heat shock

Variation in gene expression following heat shock as a result of differences in developmental temperature was assessed using procedures similar to those of Barreto et al. (2015). Offspring from 60 SD and 60 BR females were developed and acclimated at either 20 or 25°C as described above for measurements of ATP synthesis rates (with separate groups of females). In the evening prior to the last day of the adult acclimations, groups of 15 copepods (18 groups per population x treatment) were transferred to 15-mL Falcon™ conical tubes (Thermo Fisher Scientific, Waltham, MA) containing 10 mL of filtered seawater (36 ppt) with no food at the acclimation temperature of the copepods. In the morning, tubes were transferred to water baths held at 35 or 36°C for 1 h (6 per population x treatment at each temperature), and then returned to the acclimation temperature of the copepods for 1 h. The remaining 6 tubes (per population x treatment) were handled in the same manner, but remained at the acclimation temperature of the copepods for the entire 2 h. At the end of all heat shock trials, copepods were frozen at −80°C until RNA isolation.

Briefly, RNA was isolated using TRI Reagent® (Sigma-Aldrich, Inc., St. Louis, MO) with half-volume reactions according to the manufacturer’s instructions. RNA pellets were resuspended in 12 µL of Invitrogen™ UltraPure™ DNase/RNase-Free Distilled Water (Thermo Fisher Scientific, Waltham, MA) and were incubated at 56°C for 5 min. Isolations were treated with DNase using Invitrogen™ TURBO DNA-*free*™ Kits (Thermo Fisher Scientific, Waltham, MA) following the supplied protocols, and RNA concentration was determined with a NanoDrop spectrophotometer (Thermo Fisher Scientific, Waltham, MA). RNA integrity was confirmed by gel electrophoresis using two high-concentration samples. 100-150 ng of total RNA was used to synthesize cDNA for each sample with Applied Biosystems™ High-capacity RNA-to-cDNA™ Kits (Thermo Fisher Scientific, Waltham, MA) as instructed by the manufacturer, and the resulting cDNA samples were normalized to 2 ng input RNA µL^−1^.

The mRNA expression levels of heat shock protein beta 1 (*hspb1*), heat shock protein 70 (*hsp70*), mitochondrially encoded ATP synthase membrane subunit 6 (*mt-atp6*) and glyceraldehyde 3-phosphate dehydrogenase (*gapdh*) were assessed by quantitative real-time polymerase chain reaction (qRT-PCR). Primers for *hspb1*, *hsp70* and *gapdh* for the SD population were obtained from Barreto et al. (2015). If necessary due to single nucleotide polymorphisms between the populations, equivalent primers were designed for the BR population using a population-specific reference genome (Barreto et al., 2018), and *mt-atp6* primers for each population were designed using population-specific mitochondrial genomes (DQ913891; Burton et al., 2007; Barreto et al., 2018). All primer sequences are listed in Table 1. 15 µL qRT-PCR reactions were prepared in duplicate with 4 µL cDNA, 5 pmol of each primer, and 7.5 µL iTaq Universal SYBR Green Supermix (Bio-Rad Laboratories, Inc., Hercules, CA). All reactions were conducted on an AriaMx Real-time PCR System (Agilent Technologies, Inc., Santa Clara, CA) with the following protocol: 95°C for 2 min, then 95°C for 10 s followed by 58°C for 30 s for 40 cycles. The presence of a single amplicon was confirmed by a melting curve analysis after each reaction. Samples for each population were quantified relative to population-specific 5-point standard curves that were included on all reaction plates, and were prepared by serial dilution (1X to 1/625X) of a high-concentration heat shock sample from each population. Transcript levels of *hspb1*, *hsp70* and *mt-atp6* were then expressed relative to those of *gapdh*, which has been confirmed to be an appropriate housekeeping gene for heat shock studies in *T. californicus* (Schoville et al., 2012; Barreto et al., 2015; Harada and Burton, 2019). Final qRT-PCR sample sizes for the majority of our treatments and genes were n = 6; however, for some groups n = 4 or 5 due to a combination of insufficient RNA for cDNA synthesis, failed reactions as a result of extremely low *hspb1* expression levels under control conditions, or insufficient cDNA to assess all genes (one instance resulting in no estimate for *mt-atp6*). As a result, the final sample sizes for all qRT-PCR data are presented in detail in Table S2.

**Table 1.**
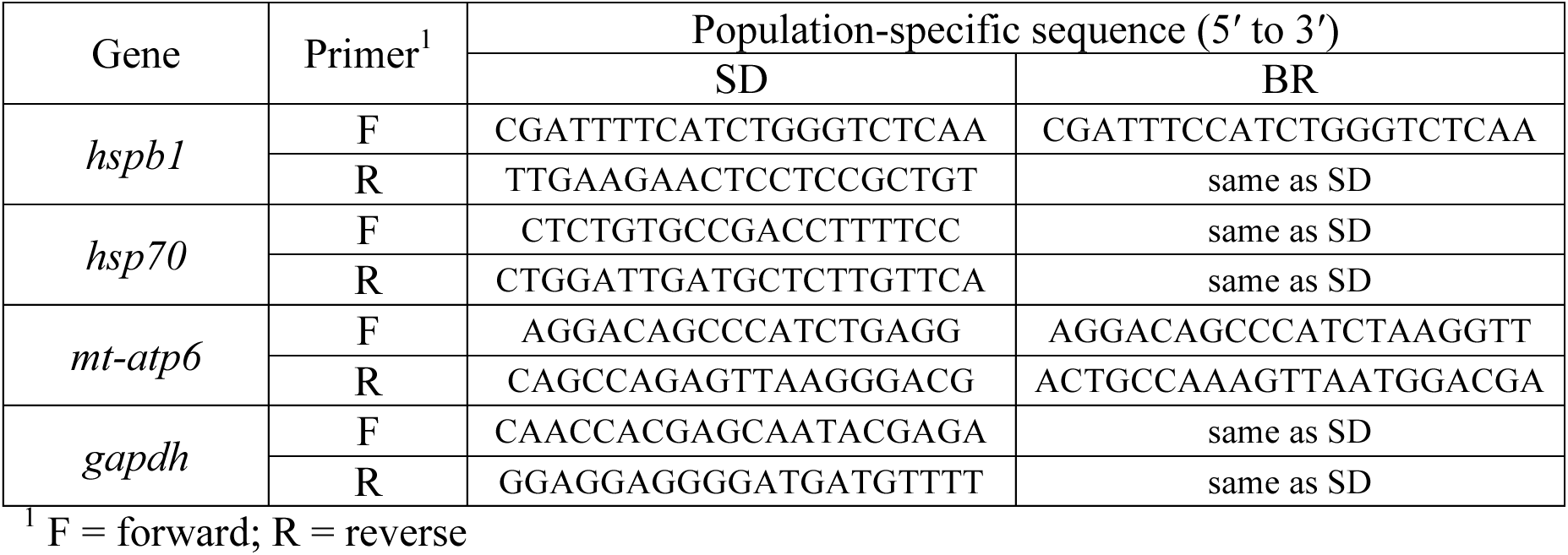
qRT-PCR primers.

### Statistical analyses

All analyses were performed with R v3.4.0 (R Core Team, 2017) and α = 0.05. Latitudinal variation in CT_max_ across all populations failed to satisfy the assumptions for parametric statistics even after log transformation. Thus, differences among populations were assessed by Kruskal-Wallis analysis of variance (ANOVA) followed by Nemenyi post-hoc tests. Potential effects of sex on CT_max_ were assessed by Wilcoxon rank-sum tests within each population. In contrast, variation in CT_max_ associated with developmental and adulthood temperatures met the assumptions for parametric tests in both the SD and BR, and the SC and PE experiments. These data were assessed by general linear models followed by ANOVAs with population, developmental temperature, and adult acclimation treatment as factors. Post-hoc comparisons among groups were performed with Tukey tests. After log transformation to meet assumptions of normality and homogeneity of variances, variation in ATP synthesis rate was assessed with a mixed-effect linear model followed by ANOVA with fixed effects of population, acclimation treatment and assay temperature, and a random effect of mitochondrial isolation. All interactions between factors were not significant in the initial model (*p* ≥ 0.12 for all), and were removed from the final model for this test. Planned pairwise post-hoc comparisons were conducted between assay temperatures within each population x acclimation treatment, between acclimation treatments within each population x assay temperature, and between populations within each acclimation treatment x assay temperature with Student’s *t* tests (84 comparisons). The resulting *p*-values were corrected for multiple tests by the Benjamini-Hochberg method (Benjamini and Hochberg, 1995). As an alternative method to examine variation in the thermal sensitivity of ATP synthesis rate, rates were normalized to the 25°C rate within each mitochondrial isolation. Variation in normalized ATP synthesis rate was assessed at assay temperatures from 30 to 37°C with similar methods to those described above for the unnormalized rates (although log transformation and removal of interactions among model factors were not required). Finally, mRNA expression data were all log transformed to meet necessary assumptions, and then differences among groups were examined by two-way ANOVAs with acclimation treatment and heat shock exposure as factors followed by post-hoc Tukey tests. Note comparisons of gene expression between populations were not made, because the expression levels of each population were quantified relative to standard curves that were population specific and for some genes the qRT-PCR primers were population-specific as well (Table 1). Full ANOVA tables for all models tested have been uploaded to the Dryad Digital Respository (*upload will be completed and the accession number provided should the manuscript be accepted*).

## Results

### Latitudinal variation in critical thermal maximum

CT_max_ demonstrated significant variation among populations (*p* < 2.2 x 10^−16^; Fig. 1), and as the geographic locations of the populations moved closer to the equator, CT_max_ generally increased, although there was limited resolution among populations in post-hoc comparisons likely due to nonparametric tests (Fig. 1). This pattern of variation in CT_max_ among populations is consistent with local thermal adaptation and with several previous studies examining latitudinal variation in upper thermal tolerance in *T. californicus* with static tolerance assays (Willett, 2010; Kelly et al., 2012; Pereira et al., 2014, 2017; Leong et al., 2018; Willett and Son, 2018; Foley et al., 2019). However, unlike previous static tolerance measurements (Willett, 2010; Willett and Son, 2018; Foley et al., 2019), CT_max_ did not vary between females and males in the current study (*p* ≥ 0.19 within all populations). In any case, these results suggest that our CT_max_ methodology effectively captures latitudinal variation in upper thermal tolerance among populations despite using relatively low numbers of individuals, which makes this approach ideal for examining effects of phenotypic plasticity on upper thermal tolerance in experiments utilizing split-clutch (i.e., divided egg sac) designs.

**Figure 1.**
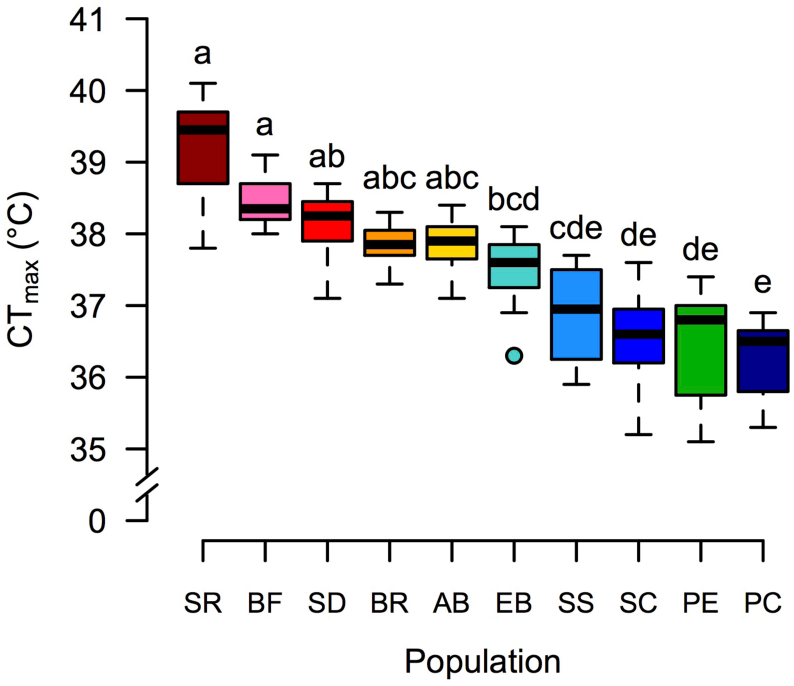
Variation in critical thermal maximum (CT_max_) among populations of *T. californicus* distributed from Mexico to Canada. Populations are plotted from southernmost to northernmost (left to right): San Rogue, Mexico (SR; dark red), La Bufadora, Mexico (BF; pink), San Diego, California (SD; red), Bird Rock, California (BR; orange), Abalone Cove, California (AB; yellow), Estero Bay, California (EB; teal), San Simeon, California (SS; light blue), Santa Cruz, California (SC; blue), Pescadero, California (PE; green) and Pacific Crest, Canada (PC; dark blue). Data are displayed as standard box and whisker plots, and lower case letters indicate the results of post-hoc comparisons among populations (n = 16 for all populations).

### Phenotypic plasticity of upper thermal tolerance

Development at 20 or 25°C resulted in variation in CT_max_ among SD and BR copepods that was significantly affected by a three-way interaction among population, developmental temperature and adult acclimation treatment (*p* = 0.03). These effects were resolved by post-hoc tests, although there was little evidence for differential effects between the populations (Fig. 2A). For all SD and BR copepods that developed at 20°C, CT_max_ values were similar regardless of adult acclimation temperature or time (range of means: 37.9 to 38.3°C; *p* = 0.054 for 20°C-developed SD acclimated to 20°C for 14 d versus 25°C for 10 d as adults, and *p* ≥ 0.22 for all other comparisons), suggesting that in 20°C-developed *T. californicus* there is no adult plasticity in upper thermal tolerance. In contrast, relative to development at 20°C, development at 25°C resulted in significant increases in CT_max_ for both SD and BR when adults were also acclimated at 25°C (range of means: 38.5 to 39.1°C; *p* ≤ 0.04 for all comparisons within populations) and these effects were similar at 10 and 14 d of acclimation (*p* ≥ 0.16 between times for both populations). These patterns are likely indicative of beneficial developmental plasticity, and match expectations given the results of a previous study utilizing static tolerance assays (Pereira et al., 2017). However, if 25°C-developed SD and BR copepods were acclimated at 20°C as adults, there was a significant loss of tolerance (i.e., decrease in CT_max_) in both populations (range of means: 37.7 to 38.1°C; *p* < 0.01 for all within population comparisons). Thus, our results are consistent with variation in adult plasticity due to differences in developmental temperature, as 20°C-developed copepods express no plasticity in adulthood, whereas 25°C-developed copepods display a clear difference in CT_max_ associated with experiencing either 20 or 25°C as adults.

**Figure 2.**
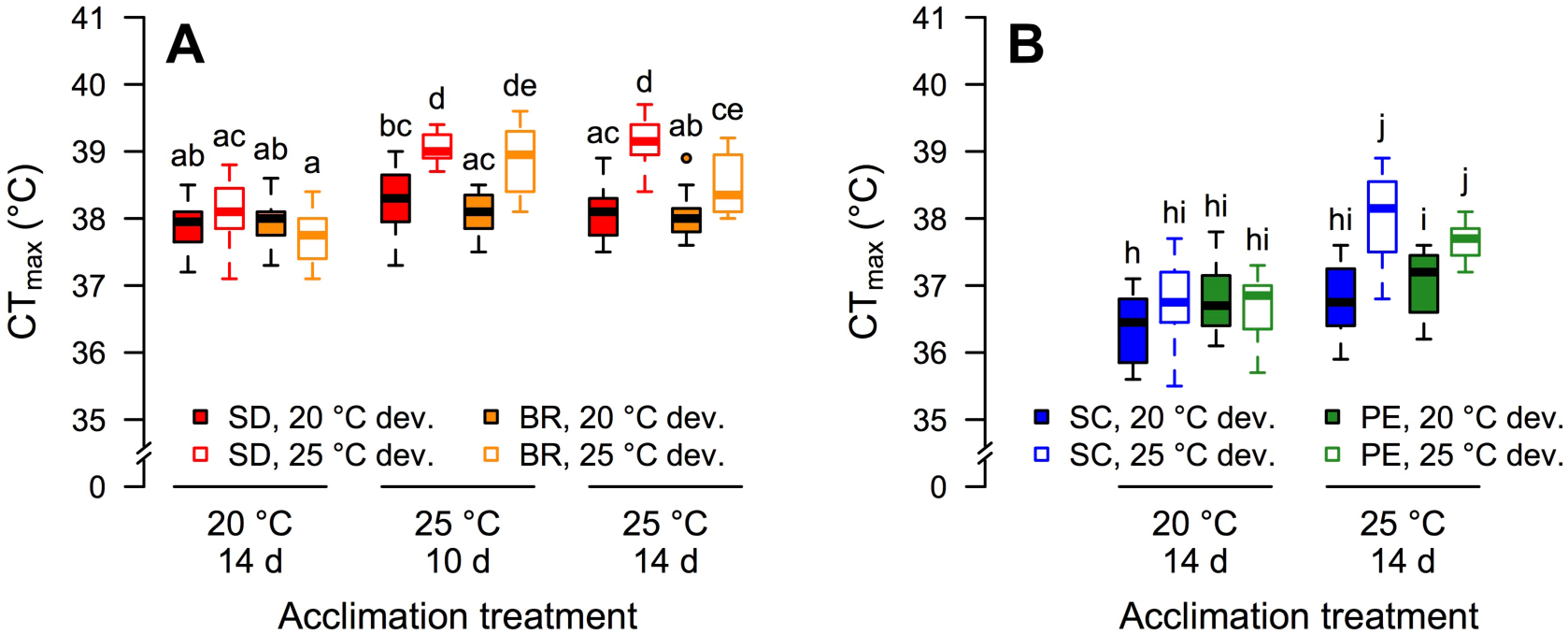
Phenotypic plasticity of critical thermal maximum (CT_max_) in Californian populations of *T. californicus* as a result of temperatures experienced during development and adulthood. Panel A: San Diego (SD; red) and Bird Rock (BR; orange) copepods. Panel B: Santa Cruz (SC; blue) and Pescadero (PE; green) copepods. Data are displayed as standard box and whisker plots (20°C development – filled boxes; 25°C development – open boxes), and lower case letters indicate the results of post-hoc comparisons among treatments within each panel (n = 16 for all groups except 25°C-developed and 25°C-acclimated PE for which n = 15).

To assess the potential for local thermal adaptation (at least along the coast of California) to shape the interaction between developmental and adult plasticity observed in SD and BR *T. californicus*, we conducted a second experiment examining plasticity of CT_max_ in the SC and PE populations, which are relatively cold-adapted (see Fig. 1). In SC and PE copepods, variation in adult CT_max_ was unaffected by the three-way interaction among population, developmental temperature and adult acclimation treatment (*p* = 0.60). Moreover, there were no significant effects of interactions between population and either developmental temperature (*p* = 0.83) or adult acclimation treatment (*p* = 0.23), or of population alone (*p* = 0.20). However, the interactive effects of developmental temperature and adult acclimation treatment significantly affected CT_max_ (*p* = 9.9 x 10^−5^). Again, post-hoc comparisons resolved this effect, and suggested similar patterns of variation among treatments for SC and PE as those described above for SD and BR (Fig. 2B). There was no significant variation in CT_max_ as a result of adult acclimation temperature in 20°C-developed copepods from either SC or PE (means: 36.4 and 36.8°C for SC, and 36.8 and 37.1°C for PE; *p* ≥ 0.39 between adult temperatures for both populations). In contrast, development at 25°C resulted in significantly higher CT_max_ in both SC and PE copepods, but only when adults were acclimated at 25°C (means: 38.0 and 37.6°C for SC and PE, respectively; *p* ≤ 0.04 for all comparisons within populations), as if 25°C-developed copepods from either population were acclimated to 20°C as adults, there was a significant decrease in CT_max_ (means: 36.7 for both SC and PE; *p* < 0.01 for both within population comparisons). Therefore, the interactive effects of developmental and adulthood temperatures in SC and PE were essentially identical to those detected for SD and BR with adult plasticity observed in 25°C-, but not in 20°C-developed copepods. Taken together, these results suggest (1) that interactive effects of plasticity at different life stages on upper thermal tolerance are likely prevalent among populations, and (2) that there is little evidence that these effects have been shaped by local thermal adaptation in the four populations examined in the current study.

### Plasticity of ATP synthesis rate

Maintaining *T. californicus* at 20 or 25°C for both development and acclimation as adults resulted in significant variation in the thermal performance curve for CI+II ATP synthesis rates in isolated mitochondria (Table 2). Specifically, these curves were affected by population (*p* = 4.4 x 10^−5^), temperature of development and acclimation (*p* < 2.2 x 10^−16^), and assay temperature (*p* < 2.2 x 10^−16^). In all developmental and acclimation treatments, synthesis rates initially increased and then decreased with increasing assay temperatures (see Table 2), and post-hoc tests found no evidence for differences between the SD and BR copepods within each treatment x assay temperature combination (*q* ≥ 0.11 for all). Yet, across all assay temperatures in both populations, 25°C development and adult acclimation resulted in higher ATP synthesis rates compared to those measured following development and acclimation at 20°C (*q* ≤ 0.046 for all). Thus, as expected based on differences in CT_max_ between these treatments (Fig. 2A), synthesis rates at high assay temperatures were greater in copepods held at 25°C than in copepods held at 20°C.

**Table 2.** Developmental and adulthood plasticity in complexes I and II (CI+II)-fueled ATP synthesis rates as a result variation in temperature.

| Population | Assay temperature (°C) | CI+II ATP synthesis rate<sup>1</sup><br>(*pmol min<sup>-1</sup> ng DNA<sup>-1</sup>*) |  |
| --- | --- | --- | --- |
|  |  | 20°C dev. & acc.<sup>2</sup> | 25°C dev. & acc.<sup>2</sup> |
| San Diego, California (SD) | 20 | 1.48 ± 0.20<sup>a</sup> | 2.58 ± 0.35<sup>h*</sup> |
|  | 25 | 1.74 ± 0.21<sup>b</sup> | 3.13 ± 0.39<sup>i*</sup> |
|  | 30 | 1.78 ± 0.22<sup>b</sup> | 3.43 ± 0.37<sup>j*</sup> |
|  | 33 | 1.42 ± 0.20<sup>a</sup> | 3.47 ± 0.41<sup>j*</sup> |
|  | 35 | 1.14 ± 0.14<sup>c</sup> | 2.88 ± 0.35<sup>i*</sup> |
|  | 37 | 0.74 ± 0.11<sup>d</sup> | 1.75 ± 0.25<sup>k*</sup> |
| Bird Rock, California (BR) | 20 | 1.34 ± 0.06<sup>a</sup> | 2.13 ± 0.26<sup>hi*</sup> |
|  | 25 | 1.58 ± 0.11<sup>b</sup> | 2.54 ± 0.34<sup>ij*</sup> |
|  | 30 | 1.56 ± 0.11<sup>b</sup> | 2.80 ± 0.41<sup>j*</sup> |
|  | 33 | 1.32 ± 0.07<sup>a</sup> | 2.47 ± 0.36<sup>i*</sup> |
|  | 35 | 0.95 ± 0.07<sup>c</sup> | 1.97 ± 0.32<sup>h*</sup> |
|  | 37 | 0.53 ± 0.03<sup>d</sup> | 1.38 ± 0.24<sup>k*</sup> |

<sup>1</sup> mean ± s.e.m.; letters indicate the results of post-hoc tests within populations and dev. & acc. treatments; asterisks indicate differences within populations and assay temperatures
<sup>2</sup> dev. & acc. = development and adult acclimation

Harada et al. (2019) specifically suggested that the capacity to maintain ATP synthesis rates at high temperatures was correlated with variation in upper thermal tolerance among populations of *T. californicus*, and this possible difference between developmental and acclimation treatments has the potential be masked by the overall vertical shift in the thermal performance curve for this trait described above (Table 2). That said, in SD copepods, rates of ATP synthesis first significantly declined with temperature between assay temperatures of 30 and 33°C for the 20°C treatment (*q* = 7.3 x 10^−4^), whereas for the 25°C treatment the first decrease occurred between assay temperatures of 33 and 35°C (*q* = 1.7 x 10^−3^), suggesting that copepods held at warmer temperatures maintained synthesis rates at higher temperatures at least in this population. As an alternative approach to examine these effects, we normalized ATP synthesis rates across temperatures to those measured at 25°C within each mitochondrial isolation for each developmental and acclimation treatment, which allows comparisons of the proportional changes in synthesis rate with temperature across treatments (Fig. 3). Note that this normalization could also reasonably be done to the rates measured at 20°C, but the results would be similar, and 25°C is a slightly more conservative choice for assessing differences at high temperature (see Fig. 3). After normalization, proportional changes in rates of ATP synthesis were affected by a three-way interaction among population, temperature of development and adult acclimation, and assay temperature (*p* = 0.02). Post-hoc comparisons revealed similar patterns of variation among assay temperatures as those detected for the unnormalized rates (as would be expected), and when assayed at 37°C 20°C-developed and -acclimated SD copepods maintained higher synthesis rates than 20°C-developed and -acclimated BR copepods (*q* = 0.03). Additionally, at high assay temperatures both populations maintained greater rates of ATP synthesis when they were held at 25°C than when they were held at 20°C (assay temperatures: 33 to 37°C for SD and 37°C for BR; *q* ≤ 5.5 x 10^−3^ for all SD and *q* = 0.02 for BR). Therefore, taken together our results suggest that development and adult acclimation at warmer temperatures in *T. californicus*, which result in increased CT_max_ (Fig. 2), also result in maintenance of ATP synthesis rates at higher temperatures.

**Figure 3.**
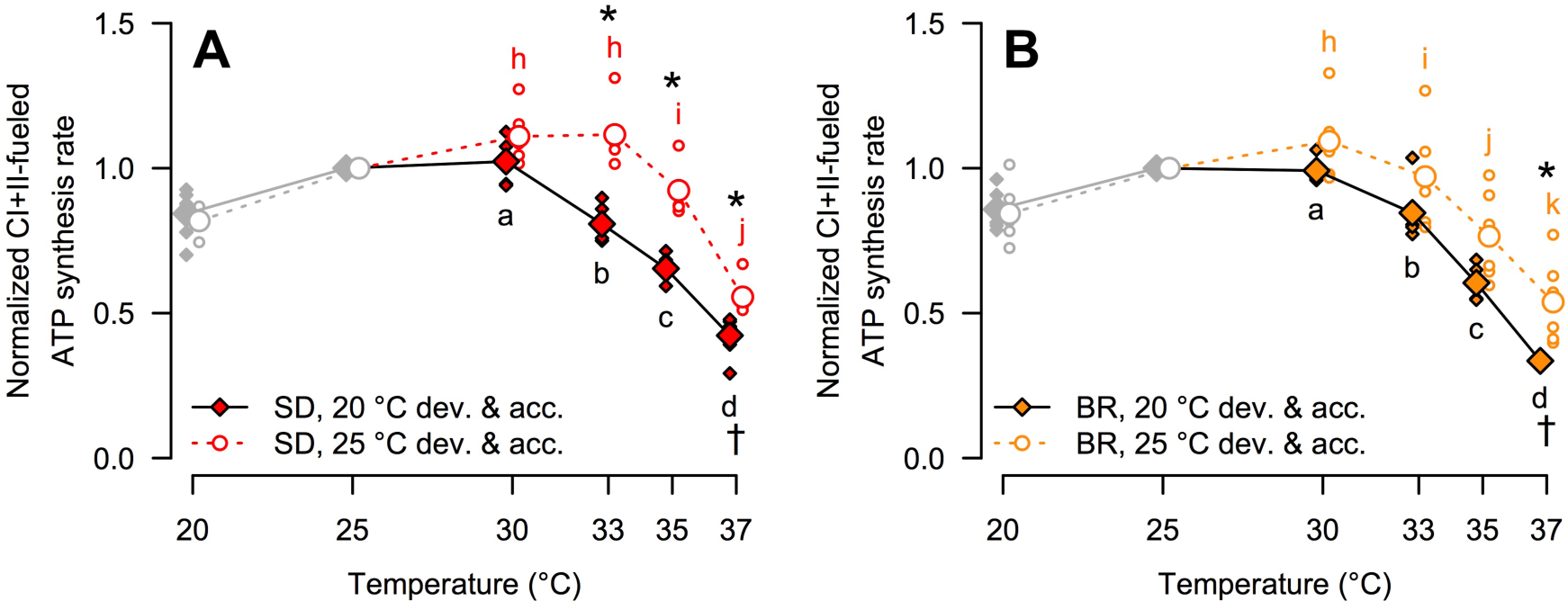
Proportional changes (from 25°C) in the thermal performance curves for complexes I and II (CI+II)-fueled ATP synthesis rates as a result of developmental and adulthood temperatures in *T. californicus*. Panel A: data for the San Diego population (SD; red), and panel B: data for the Bird Rock population (BR; orange). Copepods were developed (dev.), and then acclimated as adults (acc.) at 20°C (filled diamonds; solid lines) or 25°C (open circles; dotted lines). Small symbols display individual data points (n = 6 per population and dev. & acc. treatment), and large symbols display mean values for each group. Grey symbols show data that were not assessed statistically after normalization. Lower case letters indicate the results of post-hoc comparisons among assay temperatures within each dev. & acc. treatment for each panel, asterisks indicate differences between the dev. & acc. treatments within assay temperatures for each population, and daggers indicate differences between populations for specific assays temperatures within each dev. & acc. treatment.

### Plasticity of gene expression following heat shock

The mRNA expression levels of both heat shock proteins examined in the current study (*hspb1* and *hsp70*) demonstrated similar effects of heat shock, and developmental and adult acclimation temperature regardless of population (SD or BR; Fig. 4). In all cases, gene expression was affected by a significant interaction between the temperature of development and acclimation, and the heat shock exposure (*p* = 0.04 for *hsp70* in SD, and *p* ≤ 5.4 x 10^−3^ for all others). In general, copepods held at 25°C expressed higher levels of *hspb1* and *hsp70* than copepods held at 20°C, particularly after heat shock, although these patterns were not always resolved by post-hoc tests (see Fig. 4). However, overall these results suggest that warmer developmental and adult acclimation temperatures result in greater induction of heat shock protein expression during acute exposures to elevated temperatures in *T. californicus*, which is consistent with the difference in CT_max_ between development and acclimation at 20 or 25°C in this species (Fig. 2).

**Figure 4.**
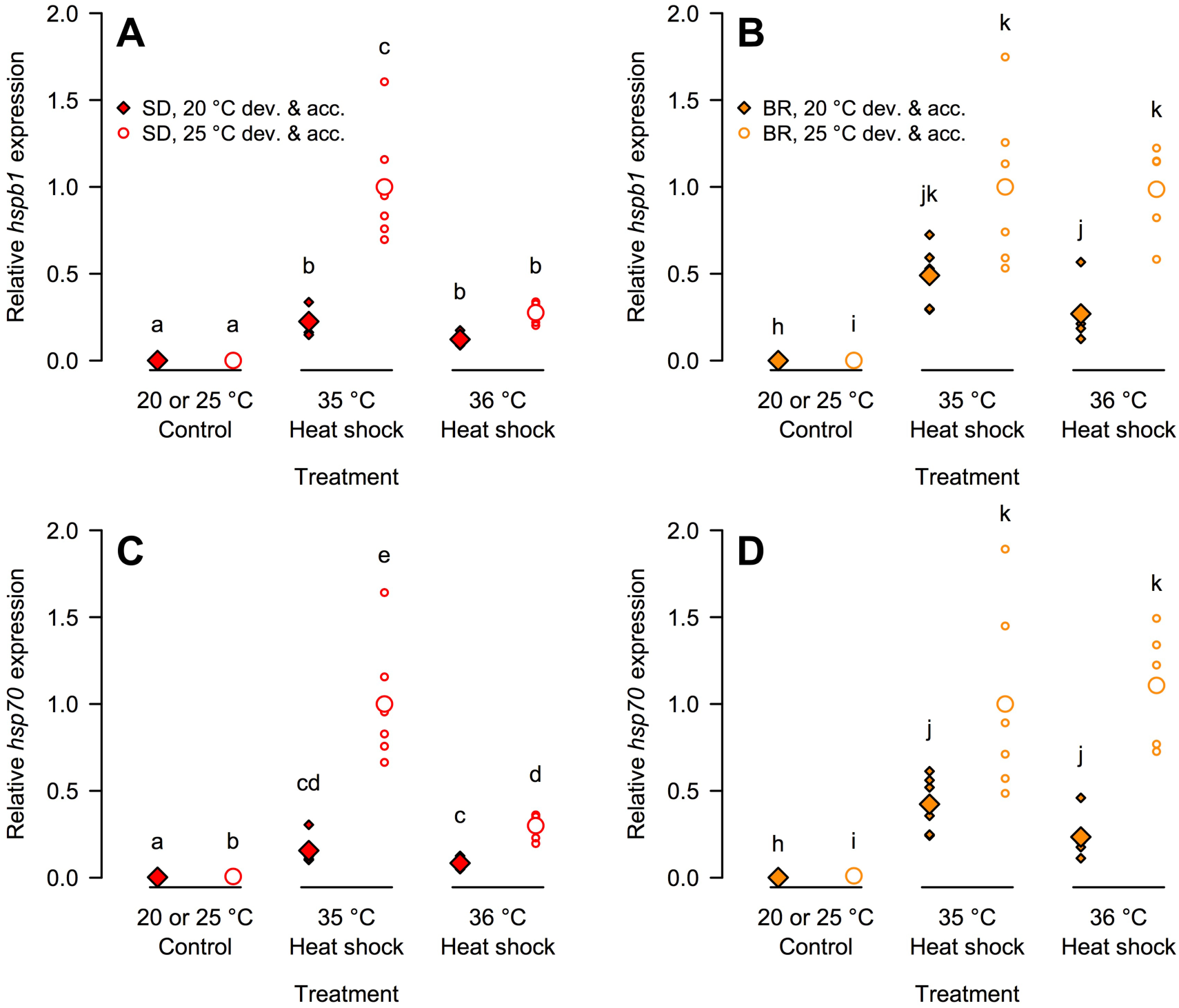
Variation in the induction of heat shock protein mRNA expression (A,B – *hspb1*; C,D – *hsp70*) as a result of developmental and adulthood temperatures in *T. californicus*. Panels A,C: data for the San Diego population (SD; red), and panels B,D: data for the Bird Rock population (BR; orange). Copepods were developed (dev.), and then acclimated as adults (acc.) at 20°C (filled diamonds; solid lines) or 25°C (open circles; dotted lines). Expression levels were quantified relative to those of the housekeeping gene *gapdh*, and are displayed normalized to the mean expression level of the 35°C heat shock treatment for the 25°C dev. & acc. copepods. Small symbols display individual data points (n = 5 or 6 for all treatments except the 36°C heat shock treatment for the SD 20°C dev. & acc. copepods for which n = 4; see Table S2 for details), large symbols display mean values for each treatment, and lower case letters indicate the results of post-hoc comparisons among treatments within each panel.

Given the potential role of mitochondrial performance in determining upper thermal tolerance (e.g., Harada et al., 2019) and a previous demonstration of decreased mitochondrial-encoded mRNA levels following heat shock in *T. californicus* (Schoville et al., 2012), we also examined variation in the expression of a mitochondrial-encoded transcript. In the current study, there were interactive effects of developmental and adult acclimation temperature, and heat shock exposure on the mRNA expression of *mt-atp6* in both the SD and BR population (*p* ≤ 0.02; Table 3). In SD copepods, *mt-atp6* levels were similar in control and 35°C-exposed animals, regardless of the temperature of development and acclimation (*p* = 1.00 for both). However, within both treatments expression levels were significantly higher in copepods developed and acclimated at 25°C than in those developed and acclimated at 20°C (*p* < 0.001 for both). In contrast, in SD copepods that had been held at 20°C, heat shock at 36°C increased *mt-atp6* expression (*p* ≤ 0.02), whereas in those that had been held at 25°C, the same exposure decreased *mt-atp6* expression (*p* < 0.001 for both). As a result, there was no effect of the temperature experienced throughout development and adulthood on *mt-atp6* mRNA levels in the 36°C heat shock treatment (*p* = 0.88). In BR copepods, variation in *mt-atp6* expression demonstrated somewhat different patterns than those observed for SD copepods. For both developmental and adult acclimation temperatures, there were trends for decreasing *mt-atp6* mRNA levels with increasing heat shock temperatures, but these patterns were only resolved in post-hoc tests in copepods developed and acclimated at 25°C between the control and the heat shock treatments (*p* < 0.001 for both; *p* ≥ 0.06 for all others). However, *mt-atp6* expression was greater in BR copepods developed and acclimated at 25°C than at 20°C in the control and in both heat shock treatments (*p* ≤ 0.02). Therefore, our data suggest (1) that heat shock has complex effects on mitochondrial-encoded transcript levels among populations of this species, and (2) that under control conditions *mt-atp6* is expressed at higher levels in *T. californicus* that are raised and held at 25°C than those that are raised and held at 20°C (Table 3).

**Table 3.** Variation in mRNA expression of *mt-atp6* associated with developmental and adulthood temperatures.

| Population | Heat shock treatment | Relative *mt-atp6* expression<sup>1</sup> |  |
| --- | --- | --- | --- |
|  |  | 20°C dev. & acc.<sup>2</sup> | 25°C dev. & acc.<sup>2</sup> |
| San Diego, California (SD) | Control | 0.28 ± 0.02<sup>a</sup> | 1.00 ± 0.03<sup>b</sup> |
|  | 35°C | 0.27 ± 0.02<sup>a</sup> | 0.98 ± 0.07<sup>b</sup> |
|  | 36°C | 0.43 ± 0.07<sup>c</sup> | 0.47 ± 0.02<sup>c</sup> |
| Bird Rock, California (BR) | Control | 0.17 ± 0.02<sup>hi</sup> | 1.00 ± 0.14<sup>j</sup> |
|  | 35°C | 0.10 ± 0.01<sup>hk</sup> | 0.28 ± 0.06<sup>i</sup> |
|  | 36°C | 0.08 ± 0.02<sup>h</sup> | 0.20 ± 0.04<sup>ik</sup> |

<sup>1</sup> mean ± s.e.m.; quantified relative to the expression of *gapdh* and normalized to expression of the 25°C dev. & acc. treatment under control conditions; letters indicate the results of post-hoc tests among groups within each population
<sup>2</sup> dev. & acc. = development and adult acclimation

## Discussion

The results presented here provide experimental support for both of our proposed hypotheses. First, variation in developmental temperature resulted in differences in the plasticity of upper thermal tolerance in adult *T. californicus*. Regardless of population, 25°C-developed copepods demonstrated clear plasticity of CT_max_ between adulthood temperatures of 20 and 25°C. In contrast, there was no evidence of plasticity in adults that had developed at 20°C. Second, differences in developmental temperatures were associated with plastic changes in two physiological mechanisms that are thought to contribute to the basis of local adaptation of upper thermal tolerance in this species, and these effects were consistent with the differences CT_max_ between the developmental treatments. Specifically, during acute exposures to high temperatures, the extents to which ATP synthesis rates were maintained and heat shock protein genes were induced were greater in copepods that were developed at 25°C than in those that developed at 20°C. Therefore, our data not only highlight the possibility for interactive effects across life stages on the plasticity of thermal tolerance in marine ectotherms, but also suggest that adaptive processes have the potential to shape these interactions due to shared underlying mechanisms, despite the lack of these effects observed among the four populations examined in the current study.

### Inter-population variation in CT_max_ is consistent with local thermal adaptation

In general, dynamic and static tolerance assays resolve similar patterns of variation among experimental groups or treatments (e.g., Ford and Beitinger, 2005; Jørgensen et al., 2019), and our previous study suggested this was also the case among three Californian populations of *T. californicus* (distributed across ∼3° latitude; Harada et al., 2019). Thus, it is not surprising that our CT_max_ measurements demonstrate substantial local thermal adaptation of upper thermal tolerance in *T. californicus*, as has been suggested in studies using static assays (Willett, 2010; Kelly et al., 2012; Pereira et al., 2014, 2017; Leong et al., 2018; Willett and Son, 2018; Foley et al., 2019). In general, thermal tolerance limits are expected to decline approximately linearly with latitude (Sunday et al., 2011); however, our results suggest this is not the case in this species. CT_max_ declines approximately linearly from San Rogue, Mexico to Estero Bay, California (−0.20°C per degree latitude), then abruptly from Estero Bay to San Simeon, California (−3.51°C per degree latitude), and then gradually from San Simeon to Pacific Crest, Canada (−0.05°C per degree latitude; Figure S1C). Leong et al. (2018) found a similar pattern in *T. californicus* with a quadratic equation resulting in the best fit between latitude and thermal tolerance among populations. It is possible this pattern reflects the effects of the California Current, which result in non-linear temperature variation along the coast particularly in the summer (Figure S1A,B). Consistent with this possibility, both Leong et al. (2018) and Pereira et al. (2017) found linear changes in upper thermal tolerance when the data were plotted against habitat temperatures. Furthermore, the change in relationship between CT_max_ and latitude in the current study occurred approximately where it might be expected (i.e., near San Simeon) if the actions of the California Current and the summer sea surface thermal cline play roles in driving inter-population variation in thermal tolerance (for example see Fig. S1B), although higher densities of sampling locations and environmental temperature measurements would be required to test this possibility in a comprehensive manner.

One distinct result of the current study was the lack of variation in CT_max_ between the sexes. There is an overall consensus that, in comparison to males, female *T. californicus* are more tolerant of stressful conditions for a wide range of abiotic factors, including temperature (Willett, 2010; Willett and Son, 2018; Foley et al., 2019). Insufficient statistical power due to nonparametric tests, and relatively low sex-specific sample sizes (n = 8) may explain the lack of sex effects in the current study. However, no population demonstrated a marginal effect of sex even if adjustments for multiple comparisons were ignored, and there were also no trends supporting differences in CT_max_ between females and males (see Fig. S1C). At least two other studies have also failed to detect differences in upper thermal tolerance between the sexes (Pereira et al., 2014, 2017). It is possible that variation in culturing conditions across studies may contribute to these results. Regardless, CT_max_ was unaffected by sex in our study, and as a result we did not consider variation between females and males further here.

### Developmental temperature and adulthood plasticity in CT_max_

Across the SD, BR, SC and PE populations of *T. californicus*, we consistently observed variation in CT_max_ plasticity in adults as a result of temperatures experienced during development. After development at 25°C, upper thermal tolerance was higher in copepods acclimated to 25°C in adulthood than in copepods acclimated to 20°C, whereas 20°C-developed copepods displayed no difference in upper thermal limits between the adult acclimation treatments. These patterns could be the result of differences in reversible adult plasticity due to developmental temperatures, or of temperature-dependent reversibility of developmental plasticity, but in either case this phenotypic variation is consistent with an interactive effects between plasticity at two life stages. To our knowledge, this is the first demonstration of interactive effects between temperatures in development and in later stages of life on thermal tolerance plasticity in a marine ectotherm. Alternatively, these patterns could be potential consequences of differences in developmental survival between 20 and 25°C as we did not directly monitor survivorship in this study; however, Pereira et al. (2017) and Harada et al. (2019) observed little evidence of differential survival at these temperatures across most *T. californicus* populations. Furthermore, previous studies have also detected interactive effects of developmental and adulthood temperatures on upper thermal tolerance in zebrafish (*Danio rerio*; Schaefer and Ryan, 2006) and fruit flies (*Drosophila sp.*; Kellermann et al., 2017; Kellermann and Sgrò, 2018), which in combination with the results of the current study suggest there is mounting evidence that interactive effects of plasticity at different life stages may be common for this trait.

In *T. californicus*, the interactive effects of plasticity at different life stages were relatively large, as in one developmental treatment (25°C) there was plasticity of CT_max_ observed in adults, whereas in the other developmental treatment (20°C) this plasticity was completely absent. Similarly, variation in developmental temperature results in presence-absence differences in adult plasticity in *D. melanogaster* (Kellermann et al., 2017), although in *Drosophila sp.* cooler developmental temperatures tend to increase adult plasticity (Kellermann et al., 2017; Kellermann and Sgrò, 2018), whereas our results suggest warmer developmental temperatures increase adult plasticity in *T. californicus*. The loss of adult CT_max_ plasticity at an only moderately reduced developmental temperature in *T. californicus* is potentially surprising given the prevalence of capacities for acclimation of this trait across many species (e.g., Gunderson and Stillman, 2015). In the current study, we examined only a relatively small range of temperatures (20-25°C), which may contribute to the lack of observed plasticity, and the extent to which plasticity may alter CT_max_ in 20°C-developed copepods over a wider range of temperatures remains an open question.

The short generation times of *T. californicus* and *Drosophila sp.* likely increase the concordance between developmental and adulthood temperatures (particularly in the intertidal habitats of *T. californicus*), which may relate to the strength of interactive effects associated with developmental temperatures as the effects of developmental plasticity are expected to be stronger if temperatures in development are predictive of those experienced as adults (Cooper et al., 2010, 2012; Nettle and Bateson, 2015; Beaman et al., 2016). Consistent with this possibility, in the comparatively long-lived zebrafish, Schaefer and Ryan (2006) observed only subtle shifts in CT_max_ plasticity in adults as a result of differences in developmental conditions. Although the interactive effects of developmental and adulthood temperatures with respect to patterns of CT_max_ plasticity were relatively strong in *T. californicus*, the maximum difference in CT_max_ among treatments was approximately 1°C, which could be interpreted as a modest change. However, in copepods held at 20°C, CT_max_ also only increased by about 1°C from the northern SC or PE populations to the southern SD or BR populations. Therefore, the magnitudes of change in upper thermal tolerance associated with 5°C variation in developmental temperature were similar to those associated with local thermal adaptation across ∼775 km of the Californian coast (∼3° latitude), implying that even a 1°C change in CT_max_ represents a relatively consequential change in upper thermal tolerance. Taken together, our data clearly demonstrate that variation in developmental and adulthood temperatures can have substantial interactive effects on the upper thermal tolerance of aquatic ectotherms.

### Mechanisms underlying CT_max_ plasticity interactions and local thermal adaptation

The possibility of interactions between adaptive processes and phenotypic plasticity is well established (e.g., Crispo, 2007; Hendry, 2016; Donelson et al., 2019; Kelly, 2019), and thus there is also the potential for local thermal adaptation to affect interactions between thermal tolerance plasticity at difference life stages. Furthermore, if heat hardening is used as a metric of plasticity in adults, there is some evidence for effects of adaptation on these interactions among *Drosophila sp.* (Kellermann and Sgrò, 2018). However, when this possibility was assessed with adult acclimations in temperate and tropical populations of *D. melanogaster*, there was little variation between populations in the patterns of phenotypic variation associated with plastic interactions (Kellermann et al., 2017). Similarly, although all of the *T. californicus* populations examined in the current study demonstrated effects of developmental temperature on adulthood plasticity, there was no variation in these interactions among populations. Indeed, the average effects of plasticity in 25°C-developed adults were remarkably similar across the four populations (SD: 1.0°C, BR: 0.8°C, SC: 1.3°C and PE 0.9°C). Thus, our results suggest that interactive effects of developmental and adulthood plasticity on CT_max_ have not been altered substantially by local thermal adaptation among these populations of *T. californicus*. However, we also found that physiological mechanisms that putatively underlie latitudinal variation in upper thermal tolerance in this species (e.g., Schoville et al., 2012; Harada et al., 2019) show patterns of variation in response to developmental temperature that parallel variation in CT_max_.

Harada et al. (2019) demonstrated that the temperatures at which maximal ATP synthesis rates in isolated mitochondria first declined with increasing temperature were correlated with differences in CT_max_ among the SD, AB and SC populations of *T. californicus*. Thus, maintaining sufficient capacity to generate ATP at high temperatures may play a role in determining upper thermal tolerance in this species. Consistent with this possibility, several studies have demonstrated loss of ATP synthesis capacity in heart mitochondria at temperatures that are approximately equal to or are immediately below the upper thermal limits in species of fishes (Iftikar and Hickey, 2013; Christen et al., 2018; O’Brien et al., 2018). Temperature-mediated plasticity of mitochondrial functions is also often observed in ectothermic species (e.g., Guderley, 2004; Seebacher et al., 2010; Chung and Schulte, 2015; Chung et al., 2017a,b, 2018; Bryant et al., 2018), and our data suggest this is the case in SD and BR copepods held at 20 or 25°C. Overall, *T. californicus* that were held at 25°C had greater ATP synthesis rates than those that were held at 20°C in both populations, which was consistent with higher expression levels of *mt-atp6* at 25°C compared to 20°C in animals under control (i.e., non-heat shocked) conditions. Additionally, 25°C copepods maintained their ATP synthesis rates to a greater extent at high temperatures than 20°C copepods, which is consistent with difference in CT_max_ between these treatments. In comparison to Harada et al. (2019), the thermal performance curves observed in our study were horizontally shifted to moderately lower temperatures, and were remarkably flat with maximum Q_10_ values of approximately 1.4-1.5 across treatments. Although relatively thermally insensitive physiological rates have been observed previously in *T. californicus* (e.g., Scheffler et al., 2019), there is clearly an unknown source of variation in these ATP synthesis curves among studies. Most likely, culturing conditions contribute to this variation, as Harada et al. (2019) examined copepods taken directly from stock cultures (i.e., 400-mL beakers), whereas in the current study we raised groups of copepods specifically for these measurements in 10-cm petri dishes. Regardless, with the exception of temperature, our 20 and 25°C treatments were held under equivalent conditions, and therefore the difference in high-temperature maintenance of ATP synthesis rates between these treatments is likely robust to any variation in thermal performance curve estimates across studies.

Schoville et al. (2012) examined genetically determined differences in the transcriptomic response to acute heat stress between the SD and SC populations of *T. californicus*. Both the strongest response and largest difference between the two populations was the extent to which heat shock protein genes were induced following heat stress. Particularly for *hspb1* and *hsp70*, heat shock protein mRNA expression was increased to much higher levels in the warm-adapted SD population than in the relatively cold-adapted SC population. As heat shock proteins are molecular chaperones that mitigate the negative effects of high temperature due to damaged and unfolded proteins (Hochachka and Somero, 2002), these transcriptomic patterns suggest that differences in heat shock protein expression may contribute to the difference in upper thermal tolerance between the SD and SC populations. The evidence for a correlation between high heat shock protein expression and increased tolerance of high temperatures is somewhat mixed among genes and species (e.g., Healy et al., 2010; Gleason and Burton, 2015); however, studies in fruit flies (*D. melanogaster*) and marine snails (*Chlorostoma funebralis*) generally support a positive relationship between these two traits (e.g., Bettencourt et al., 2008; Tomanek et al., 2008). In *T. californicus*, Tangwancharoen et al. (2018) demonstrated putatively adaptive functional variation associated with sequence differences among populations in both the regulatory and coding regions of *hspb1*, and Barreto et al. (2015) utilized RNA interference to show that knockdown of transcripts for this gene directly decreases survivorship following acute thermal stress. Therefore, the increased inductions of *hspb1* and *hsp70* we observed in SD and BR copepods held at 25°C compared to those held at 20°C are likely beneficial effects of plasticity, and are consistent with tolerance differences between these treatments. Part of this variation in heat shock protein expression may be associated with the differences in recovery temperatures in our study, as each developmental and acclimation treatment was recovered at its holding temperature. However, in all cases, the fold difference in expression between the 20 and 25°C copepods is greater than would be expected due to thermodynamic effects on reaction rates alone. Taken together, our mechanistic results indicate that both the extents to which ATP synthesis rates are maintained and to which heat shock proteins are induced at high temperatures are elevated in *T. californicus* that are developed and acclimated at 25°C compared to those that are developed and acclimated at 20°C. Thus, these mechanisms that likely contribute to local thermal adaptation of upper thermal tolerance in this species also may play a role in plastic effects due to developmental temperatures.

## Conclusion

The effects of environmental change on organisms ultimately depend on the capacities to modulate key physiological traits to facilitate performance and persistence. Phenotypic plasticity, adaptation and interactions between these processes all play important roles in these responses (e.g., Kellermann and van Heerwaarden, 2019). Here we show that interactive effects of phenotypic plasticity across life stages in the intertidal copepod *T. californicus* also contribute to variation in upper thermal tolerance. These effects may be particularly relevant for aquatic ectotherms as thermal tolerance limits likely underlie geographic range limits in many of these species (Sunday et al., 2012; Pinsky et al., 2019). Our results suggest that beneficial effects of developmental plasticity with respect to environmental change have the potential to be overestimated if considered without accounting for temperature variation in adulthood. Additionally, the data presented here indicate that the mechanisms underlying these interactive effects (e.g., shifts in the thermal performance curve for ATP synthesis and the regulation of heat shock genes) are, at least to some extent, shared with the mechanisms associated with local thermal adaptation in *T. californicus*. This mechanistic overlap highlights the potential for interactions among local adaptation and plasticity at difference life stages to shape variation in upper thermal tolerance in ectothermic organisms.

## List of abbreviations

ANOVA: Analysis of variance
CT_max_: Critical thermal maximum
CI+II: Electron transport system complexes I and II
qRT-PCR: Quantitative real-time polymerase chain reaction
Q_10_: Temperature coefficient

## Competing interests

No competing interests declared

## Funding

This work was supported by the National Science Foundation (NSF) [DEB1551466 to R.S.B.].

## Data availability

All data collected for the current study have been uploaded to the Dryad Digital Repository (*upload will be completed and accession number provided should the manuscript be accepted*).

